# SETD1B controls cognitive function via cell type specific regulation of neuronal identity genes

**DOI:** 10.1101/2020.08.07.240853

**Authors:** Alexandra Michurina, Sadman Sakib, Cemil Kerimoglu, Dennis Manfred Krüger, Lalit Kaurani, Rezaul Islam, Tonatiuh Pena Centeno, Julia Cha, Xingbo Xu, Elisabeth M. Zeisberg, Andrea Kranz, Francis Adrian Stewart, Andre Fischer

## Abstract

Histone-3-lysine-4-methylation (H3K4me) is mediated by six different lysine methyltransferases (KMTs). Amongst these enzymes SET domain containing 1b (SETD1B) has been linked to intellectual disability but its role in the adult brain has not been studied yet. Here we show that mice lacking *Setd1b* from excitatory neurons of the adult forebrain exhibit severe memory impairment. By combining neuron-specific ChIP-seq, RNA-seq and single cell RNA-seq approaches we show that *Setd1b* controls the expression of neuronal-identity genes with a broad H3K4me3 peak linked to learning and memory processes. Our data furthermore suggest that basal neuronal gene-expression is ensured by other H3K4 KMTs such as *Kmt2a* and *Kmt2b* while the additional presence of *Setd1b* at the single cell level provides transcriptional consistency to the expression of genes important for learning & memory.

## INTRODUCTION

Cognitive diseases are a heterogeneous group of disorders that depend on complex interactions of genetic and environmental factors that activate epigenetic processes(Fischer, 2014). In addition, mutations in genes that control epigenetic gene-regulation are over-represented in cognitive diseases (Kleefstra *et al*, 2014). Therefore, targeting the epigenome has emerged as a promising therapeutic avenue to treat neurodegenerative and neuropsychiatric diseases (Nestler *et al*, 2015) (Fischer, 2014). To understand the regulation of epigenetic gene-expression in the adult brain is thus of utmost importance. Histone 3 lysine 4 methylation (H3K4) is enriched around transcription start site (TSS) regions of actively transcribed genes when trimethylated (H3K4me3) (Guenther *et al*, 2006). In the human brain reduced H3K4me3 has been observed in cognitive diseases such as autism spectrum disorder (Shulha *et al*, 2012)or Alzheimer’s disease (Gjoneska *et al*, 2015a) (Kerimoglu *et al*, 2017a). In mammals, H3K4 methylation is mediated by six different lysine-methlytransferases (KMT’s), namely KMT2A (Mll1), KMT2B (Mll2), KMT2C (Mll3), KMT2D (Mll4), SETD1A, and SETD1B that catalyze mono-, di- and trimethylation (Shilatifard, 2012). The role of these enzymes in the adult brain is only beginning to emerge. Recent reports showed that *Kmt2a* and *Kmt2b* are required for hippocampus-dependent memory formation (Gupta *et al*, 2010) (Kerimoglu *et al*, 2013) (Kerimoglu *et al*, 2017b), while *Setd1a* has been linked to schizophrenia (Mukai *et al*, 2019; Singh T *et al*, 2016; Takata *et al*, 2016). *Setd1b* has been studied during development (Brici *et al*, 2017; Schmidt *et al*, 2018). Virtually nothing is known about the function of *Setd1b* in the adult brain, although mutations in *Setdb1* have been linked to intellectual disability (Hiraide *et al*, 2018; Labonne *et al*, 2016). To elucidate the role of *Setd1b* in the brain we generated mice that lack *Setd1b* from excitatory neurons of the adult forebrain. Our data reveal that *Sed1b* is essential for memory formation. Moreover, we provide evidence that *Setd1b* controls the expression of neuronal-identity genes that are characterized by a broad H3K4 trimethylation peak at the TSS, high expression levels and are intimately linked to learning and memory processes. Comparison of our data to those from other H3K4 KMTs suggest that this role is specific to *Setd1b* which provides to neurons transcriptional consistency to the expression of learning and memory genes.

## RESULTS

### Loss of Setd1b in adult forebrain neurons impairs hippocampus-dependent memory formation

To study the role of *Setd1b* in the adult brain, we crossed mice in which exon 5 of the *Setd1b* gene is flanked by loxP sites to mice that express CRE-recombinase under control of the CamKII promoter. This approach ensures deletion of *Setd1b* from excitatory forebrain neurons of the adult brain (cKO mice). Quantitative PCR (qPCR) analysis confirmed decreased expression of Setd1b from the hippocampal Cornu Ammonis (CA) area, the dentate gyrus (DG) and the cortex when compared to corresponding control littermates that carry loxP sites but do not express CRE recombinase (control group). Expression in the cerebellum was not affected confirming the specificity of the approach **(Fig 1A).** Residual expression of *Setd1b* is most likely due to the fact that deletion is restricted to excitatory neurons while other cell types are unaffected. In line with the qPCR data, SETD1B protein levels were reduced in the hippocampal CA region of *Setd1b* cKO mice **(Fig 1B)**. *Setd1b* cKO mice did not show any gross abnormalities in brain anatomy as evidenced by immunohistological analysis of DAPI staining, staining of marker-proteins for neuronal integrity Neuronal N (NEUN), microtubule-associated protein 2 (MAP2) as well as ionized calcium-binding adapter molecule 1 (IBA1) as a marker for microglia and glial fibrillary acidic protein (GFAP) as a marker for astrocytes **(Fig. 1C)**. Next, we subjected *Setd1b* cKO and control mice to behavior testing. Notably, it was previously shown that heterozygous mice expressing CRE under control of the CamKII promoter do not differ from wild type littermates (Kuczera *et al*, 2010) (Stilling *et al*, 2014) and we have confirmed this in the context of the present study also for behavior testing (**Fig. S1)**. There was no difference amongst groups in the open field test, suggesting that explorative behavior is normal in *Setd1b* cKO mice **(Fig 1D)**. Short term memory was assayed via the T-maze and was also similar amongst groups **(Fig 1E)**. Next, we subjected mice to the Morris Water Maze test to study hippocampus-dependent spatial memory. While control mice were able to learn the task as indicated by a reduced escape latency throughout the 10 days of training, *Setd1b* cKO mice were severely impaired **(Fig 1F)**. We also performed a more sensitive analysis using a modified version of the MUST-C algorithm to measure the different spatial strategies that represent either hippocampus-dependent or independent abilities (Illouz *et al*, 2016). Our results indicate that *Setd1b* cKO mice fail to adapt hippocampus-dependent search strategies such as “direct”, “corrected” and “short-chaining” **(Fig 1G)**. Consistently, the cumulative learning score calculated on the basis of these search strategies was severely impaired in *Setd1b* cKO mice **(Fig 1H)**. To assess memory retrieval, a probe test was performed. *Set1b* cKO mice were severely impaired during the probe test performed at the end of the training **(Fig 1I)**. These data show that deletion of *Setd1b* from excitatory neurons of the adult forebrain leads to severe impairment of hippocampus-dependent learning and memory abilities.

**Figure 1.**
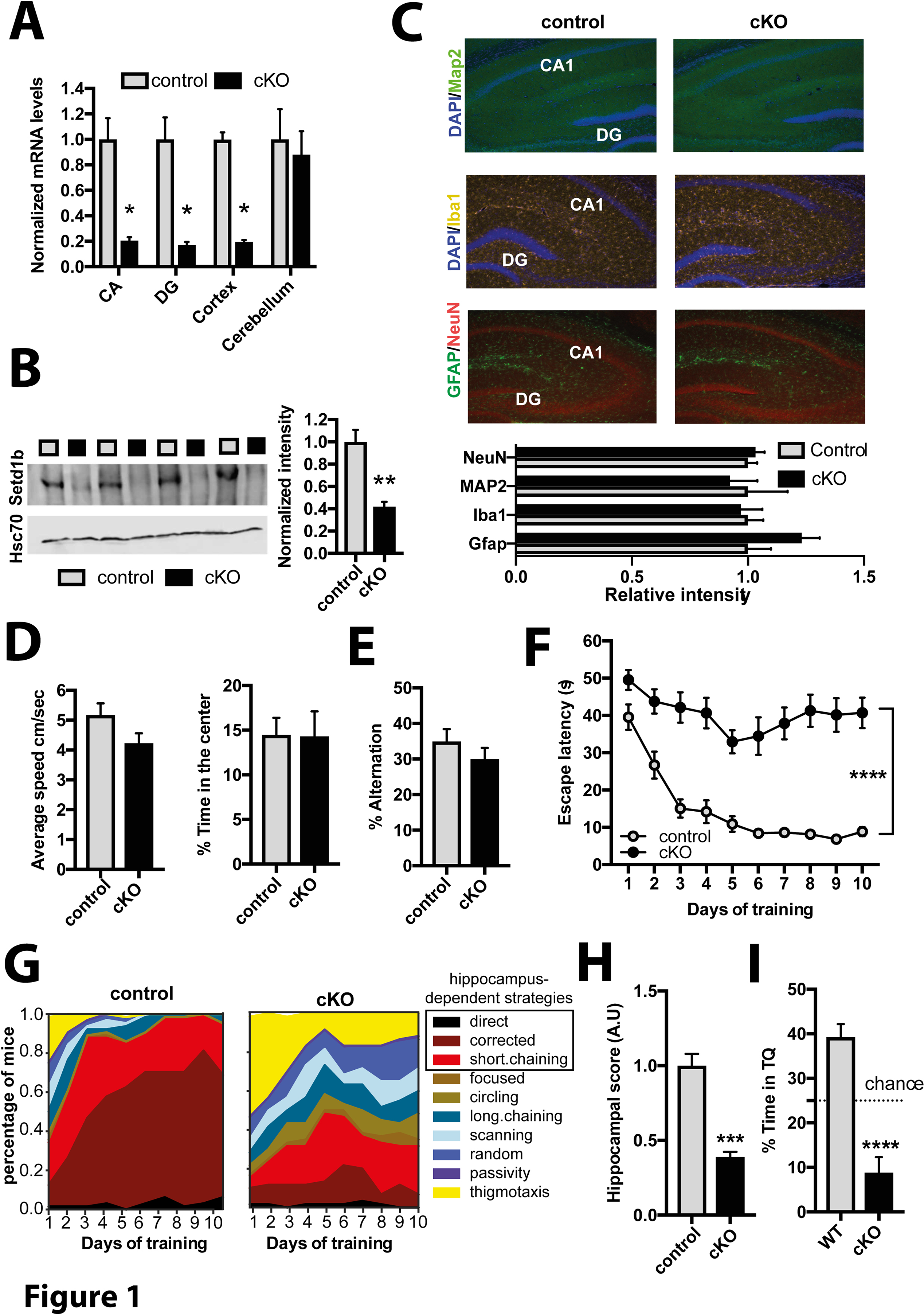
Setd1b is required for hippocampus-dependent memory. **A.** qPCR analysis shows loss of *Setd1b* in forebrain regions while levels in the cerebellum are not affected (CA: Control, n = 6; cKO, n = 6. DG: Control, n = 6; cKO, n = 6. Cortex: Control, n = 7; cKO, n = 7. Cerebellum: Control, n = 4; cKO, n = 4). * *p*-value < 0.05 (Student t-test). **B.** Immunoblot analysis shows loss of SETD1B in the hippocampus of *Setd1b* cKO mice (Control, n = 4; cKO, n = 4). ** *p*-value < 0.01 (Student t-test). **C.** Immunohistochemical staining (upper level) for marker proteins of neuronal integrity and quantification (lower panel) shows no difference in control and *Setd1b* cKO mice (NeuN: Control, n = 6; cKO, n = 6; Student t-test *p*-value = 0.57. MAP2: Control, n = 4; cKO, n = 4; Student t-test *p*-value = 0.72. Iba1: Control, n = 5; cKO, n = 5; Student t-test *p*-value = 0.8. Gfap: Control, n = 5; cKO, n = 5; Student t-test *p*-value = 0.09.). Scale bar: 100 μm. **D.** Average speed (left panel) and time spent in the center (right) panel during exposure to the open field test was similar in control and *Setd1b* cKO mice (Average speed: Control, n = 15; cKO, n = 15; Student t-test *p*-value = 0.075. Time spent in center: Control, n = 15; cKO, n = 15; Student t-test *p*-value = 0.96). **E.** Short term memory was not affected in control and *Setd1b* cKO mice as indicated by similar percent of alternations in the Y-maze test (Control, n = 15; cKO, n = 15; Student t-test *p*-value = 0.3). **F.** Escape latency during water maze training indicated severe learning impairment in *Setd1b* cKO mice (Control: n = 15, cKO: n = 15. Repeated measures ANOVA, genotype effect: F (1,28) = 82.34, **** *p*-value < 0.0001). **G.** Plots showing the specific search strategies during water maze training. Note the failure of *Setd1b* cKO mice to adapt hippocampus-dependent search strategies. **H.** The cognitive score calculated on the basis of the hippocampal search strategies is severely impaired in *Setd1b* cKO mice (Student t-test: *** p-value < 0.001). **I.** Time spent in the target quadrant during the probe test is impaired in *Setd1b* cKO mice (Control: n = 15, cKO: n = 15. **** Student t-test < 0.0001). Error bars indicate SEM.

### Setd1d controls neuronal H3K4 methylation

To elucidate the molecular mechanisms by which *Setd1b* contributes to memory formation we decided to test its impact on epigenetic gene-expression in hippocampal neurons. To this end we isolated the hippocampal CA region from *Setd1b* cKO and control mice and prepared nuclei using modified fixation protocols that allowed us to perform neuron-specific chromatin-immunoprecipitation (ChIP) to study histone-modifications and RNA-sequencing to assay gene-expression from the same samples **(Fig 2A, Fig S2)**. Since SETD1B is a histone 3 lysine 4 (H3K4) methyltransferase we decided to analyze tri-methylation (H3K4me3) of histone 3 lysine 4 that is enriched at the transcription start site (TSS) of active genes and is associated with euchromatin and active gene-expression. H3K4 methylation is believed to be a stepwise process and recent data suggest that the different methylation states (from mono- to tri-methylation) at the TSS of a gene form a gradient reflecting its specific transcriptional state (Choudhury *et al*, 2019; Soares *et al*, 2017). Thus, we also analyzed mono-methylation of histone 3 at lysine 4 (H3K4me1). In addition, we analyzed histone 3 lysine 9 acetylation (H3K9ac), an eu-chromatin mark that was shown to partially depend on H3K4 methylation (Kerimoglu *et al.*, 2013; Kerimoglu *et al.*, 2017b). Finally, we also performed Chip-seq for histone 3 lysine 27 acetylation (H3K27ac), another euchromatin mark that is linked to active gene-expression and marks promoter elements around the TSS but also enhancer regions and has not be directly linked to H3K4me3 in brain tissue. We observed that loss of *Setd1b* leads to a substantial decrease in neuronal H3K4me3 across the genome while the majority of significant changes are localized to regions in close proximity to the transcriptional start site (TSS) **(Fig 2B, C)**. Similar changes were observed for neuronal H3K9ac and H3K27ac, although less regions were affected when compared to H3K4me3 (**Fig. S3)**. We also observed significantly altered H3K4me1 in neurons of *Setd1b* cKO mice **(Fig 2B).**These changes were also almost exclusively detected in vicinity to the TSS **(Fig 2B)** but in contrast to the other investigated histone-modifications, many of the significantly altered genomic regions exhibited increased H3K4me1 levels in *Setd1b* cKO mice **(Fig. 2B, C)**. To further analyze these data, we first asked if the observed changes in histone-modifications occur within the same genomic regions. As expected the number of genes showing reduced H3K4me3 exceeded by far the number of genes showing reduced levels of H3K9ac, H3K27ac or altered H3K4me1 (Fig S3). Nevertheless, almost all regions exhibiting decreased H3K9ac where also marked by decreased H3K4me3, while the regions showing decreased H3K27ac were mainly localized to different genes (**Fig. S3**). These data support previous findings, showing that H3K4me3 is functionally linked to H3K9ac (Kerimoglu *et al.*, 2013; Kerimoglu *et al.*, 2017b) and suggest that the observed changes in H3K27ac are mainly due to secondary effects. Interestingly, decreased H3K4me3 in *Setd1b* cKO manifested exclusively downstream of the TSS, indicating that loss of *Sedt1b* may affect peak width **(Fig 2D)**. We decided to further explore this observation and noticed that there was an obvious difference amongst the genes that exhibit decreased H3K4me3 and increased H3K4me1 **(Fig. 2E)** when compared to genes that show exclusively decreased H3K4 methylation around the TSS **(Fig 2F)**. Namely, the change in H3K4me3 was most significant in genes with decreased H3K4me3 and increased H3K4me1 and was characterized by a substantially reduced H3K4me3 peak width **(Fig 2E)**, when compared to genes with decreased H3K4me3 and H3K4me1 **(Fig 2F)**. Findings from other cell types suggest a gradient of H3K4 methylation states in which the proximity of the mark to the TSS is correlated to the level of gene-expression. Thus, genes with broader H3K4me3 peaks at the TSS exhibit the highest and most consistent expression levels and represent genes of particular importance for cellular identity (Benayoun *et al*, 2015; Soares *et al.*, 2017). Indeed, our data revealed that the genes which are characterized by decreased H3K4me3 and increased H3K4me1 in *Setd1b* cKO mice, already exhibit significantly broader H3K4me3 peaks under basal conditions, when compared to genes characterized by decreased H3K4me3 but either decreased or unchanged H3K4me1 levels **(Fig 2 G)**. Interestingly, these genes were also expressed at significantly higher levels under baseline conditions **(Fig 2 H)**. Taken together, our findings suggest that *Setd1b* may be of particular importance for the expression of genes linked to the specific function of hippocampal neurons. In line with this, functional pathway analysis revealed that the genes with decreased H3K4me3 and increased H3K4me1 and thus having the broadest H3K4me3 peak under basal conditions, represent pathways intimately linked to the function of excitatory hippocampal neurons **(Fig 2I)**. Most importantly, this was not the case for the genes of the other two categories **(Fig 2I).** In summary, our data show that loss of *Setd1b* from hippocampal neurons leads to distinct changes in neuronal histone-methylation and point to a specific role of *Setd1b* in the expression of genes essential for neuronal identity and cognitive function.

**Figure 2.**
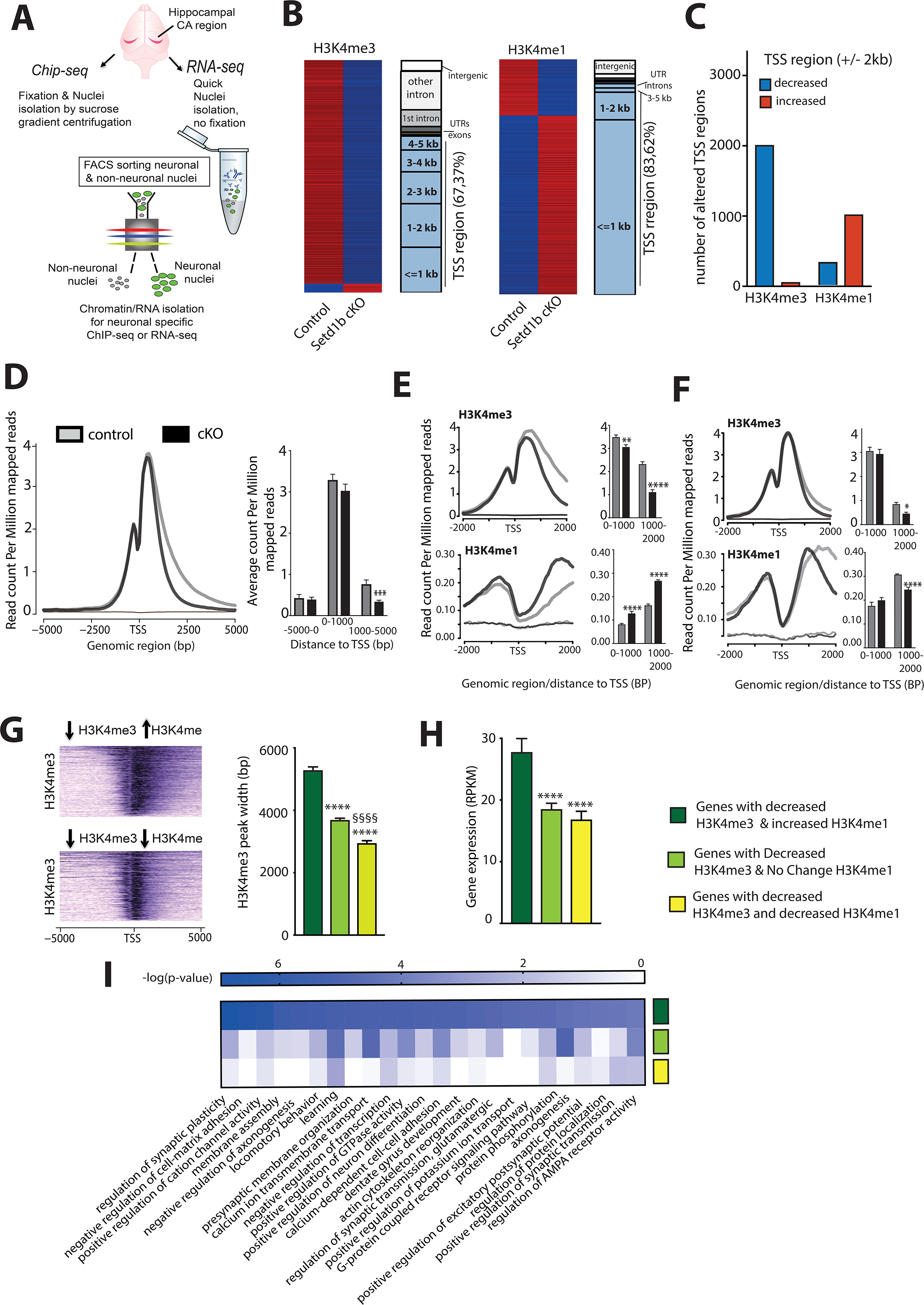
*Setd1b* controls histone-methylation and H3K4me3 peak width. **A.** Experimental scheme showing our approach to perform cell-type specific ChIP-seq and RNA-seq. For Chip-Seq we employed n = 4 for control and n = 4 form *Setd1b* cKO. **B.** Left panel: Heat map showing genes with significantly differing H3K4me3 sites at the TSS (+/− 2kb) in *Setd1b* cKO mice and the overall genomic locations of altered H3K4me3 levels. Right panel shows the same analysis for H3K4me1 (FDR < 0.05 & |fold change| > 1.5). **C.** Bar plot showing the number of genes with decreased and increased H3K4me3 and H3K4me1 at the TSS region in *Setd1b* cKO mice (FDR < 0.05 & |fold change| > 1.5). **D.** NGS plot showing H3K4me3 across all genes with significantly reduced H3K4me3 in *Setd1b* cKO mice. Left panel shows a bar chart indicating that reduced H3K4me3 in *Setd1b* cKO mice is mainly occurring downstream of the TSS (*** Student t-test *p*-value < 0.001). **E.** NGS plots showing the distribution of H3K4me3 and H3K4me1 at the close vicinity of TSS of genes that show significantly reduced H3K4me3 and increased H3K4me1 in *Setd1b* cKO mice. Bar graphs on the left show corresponding quantification (Student t-test: ** *p*-value < 0.01, **** *p*-value < 0.0001). **F.** NGS plot showing the distribution of H3K4me3 and H3K4me1 at the TSS of genes that show both reduced H3K4me3 and H3K4me1 in *Setd1b* cKO mice. Bar graphs on the left show corresponding quantification (Student t-test: * *p*-value < 0.05, **** *p*-value < 0.0001). **G.** Heatmap (left panel) showing basal state H3K4me3 peak width for genes characterized by decreased H3K4me3 in combination with either increased or decreased H3K4me1 in *Setd1b* cKO mice. Right panel: Quantification of the peak width in genes with decreased H3K4me3 in combination with either increased, decreased or not altered H3K4me1 in *Setd1b* cKO mice (One-way ANOVA: *p*-value < 0.0001. Post-hoc multiple comparisons, Tukey’s test: increased H3K4me1 vs no change H3K4me1, **** *p*-value < 0.0001; increased H3K4me1 vs decreased H3K4me1, **** *p*-value < 0.0001; no change H3K4me1 vs decreased H3K4me1, §§§§ *p*-value < 0.0001). **H.** Bar graph showing the basal wild type expression level for the 3 categories of genes that display altered H3K4me3 in *Setd1b* cKO mice. Please note that basal expression level is highest for genes with decreased H3K4me3 in combination with increased H3K4me1 that are characterized by broad H3K4me3 peaks (One-way ANOVA: *p*-value < 0.0001. Post-hoc multiple comparisons, Tukey’s test: increased H3K4me1 vs no change H3K4me1, **** *p*-value < 0.0001; increased H3K4me1 vs decreased H3K4me1, **** *p*-value < 0.0001; no change H3K4me1 vs decreased H3K4me1, *p*-value = 0.6967). **I.** Heat map showing functional pathways for the 3 categories of genes affected by reduced H3K4me3 in *Setd1b* cKO mice. Error bars indicate SEM.

### Setd1b controls the levels of highly expressed neuronal genes characterized by a broad H3K4me3 peak at the TSS

To test the impact of *Setd1b* on gene-expression directly, we analyzed the RNA-sequencing data obtained from neuronal nuclei of the same hippocampi used to generate ChIP-seq data (See Fig 2A, Fig S2). In line with the established role of H3K4me3 in active gene-expression, we mainly detected down-regulated genes when comparing control to *Setd1b* cKO mice **(Fig 3A)**. In fact, the comparatively few up-regulated genes were all lowly expressed at baseline conditions suggesting rather unspecific effects (RPKM down-regulated genes =18.77 +/−1.45 vs. up-regulated genes RPKM = 3.08 +/− 0.26; *P* < 0.0001). Further analysis revealed that the TSS of genes down-regulated in *Setd1b* cKO mice is characterized by significantly reduced H3K4me3 peak-width and increased H3K4me1 **(Fig 3B, C)**. This observation was specific to the genes down-regulated in *Setd1b* cKO mice, since random sets of genes that were not de-regulated in *Setd1b* cKO mice show normal H3K4me3 and H3K4me1 levels at the TSS **(Fig 3D)**. We also observed that the genes down-regulated as a result of *Setd1b* deletion were characterized by a significantly broader H3K4me3 peak and higher expression under basal conditions **(Fig 3C, E)**. A functional pathway analysis revealed that the genes down-regulated in *Setd1b* cKO mice are intimately linked to synaptic plasticity and learning and memory related processes **(Fig 3F)**. Taken together, these data further suggest that *Setd1b* controls a specific set of genes that are characterized by a broad H3K4me3 peak at the TSS, are highly expressed in hippocampal neurons under basal conditions and play a specific role in neuronal plasticity and identity.

**Figure 3.**
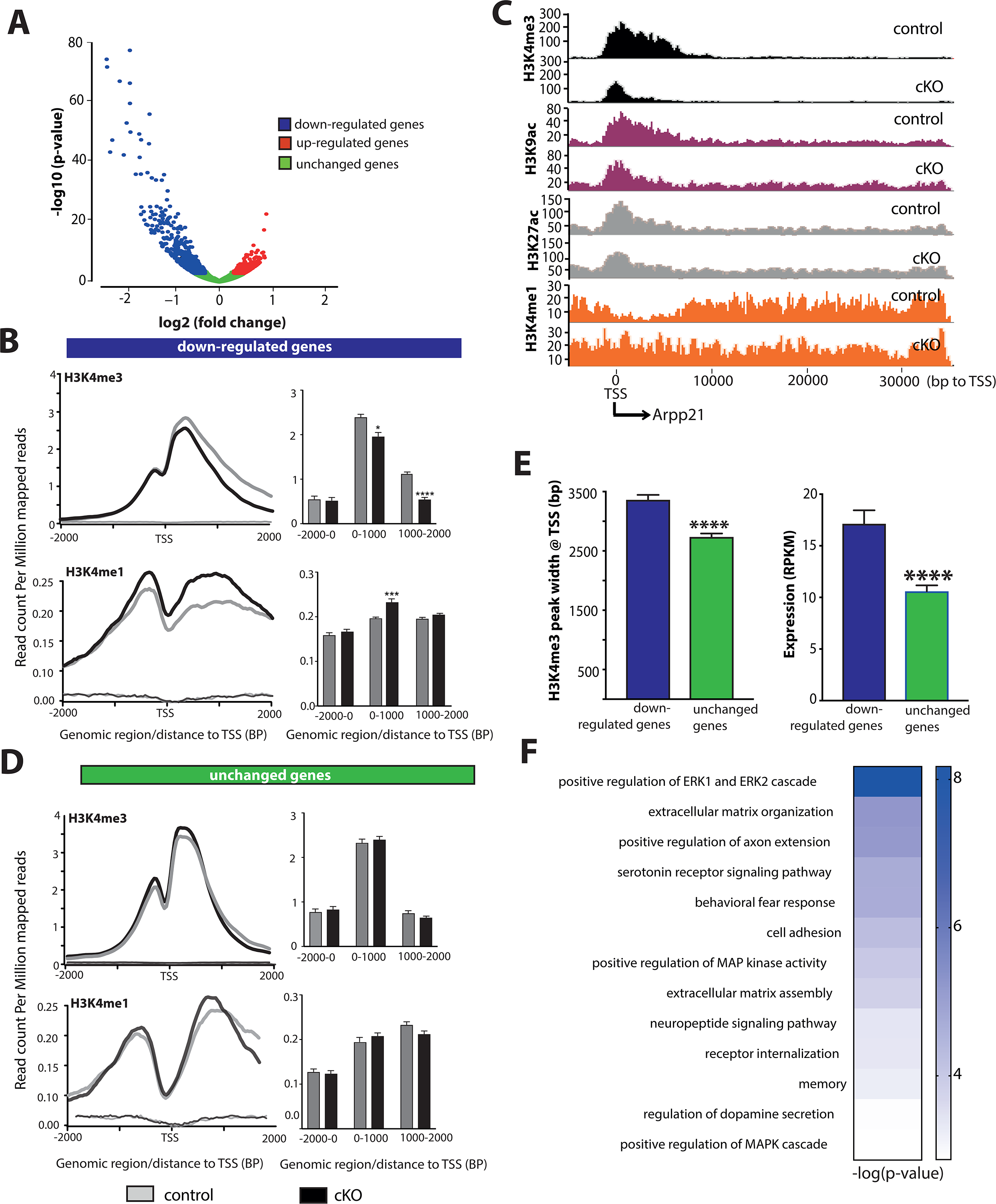
Hippocampal *Setd1b* controls highly expressed learning and memory genes characterized by a broad H3K4me3 peak. **A.** Volcano plot showing genes differentially expressed in hippocampal neurons of *Setd1b* cKO mice. n = 3/group. **B.** NGS plots showing H3K4me3 and H3K4me1 at the TSS of genes down-regulated in *Setd1b* cKO mice. Bar plots (right panel) show quantification. **C.** *Arpp21* (CAMP Regulated Phosphoprotein 21) was selected as a representative gene down-regulated in hippocampal neurons of *Setd1b* cKO mice to illustrate changes of the analyzed histone-modifications. Please note that the H3K4me3 peak-width is substantially shrinking in *Setd1b* cKO mice. At the same time there is an obvious increase of H3K4me1 at the TSS of *Arpp21* in *Setd1b* cKO mice. **D.** NGS plots showing H3K4me3 and H3K4me1 at the TSS of a random set of genes that were not altered in *Setd1b* cKO mice. Bar plot (right panel) show quantification. **E.** Left panel: H3K4me3 peaks are significantly broader in genes that are down-regulated in *Setd1b* cKO mice, when compared to a random set of genes that were unaffected. Right panel: Genes down-regulated in *Setd1b* cKO mice are characterized by higher baseline expression when compared to a random set of genes that were unaffected. **F.** Heat map showing functional pathways affected by genes down-regulated in *Sed1b* cKO mice. Error bars indicate SEM. Student t-test: * *p*-value < 0.05, **** *p*-value < 0.0001

### The regulation of highly expressed neuronal identity genes with broad H3K4me3 peaks is a specific feature of Setd1b

To provide further evidence for the specific role of *Setd1b* in the regulation of neuronal plasticity and neuronal identity genes we decided to compare *Setd1b* to other mammalian H3K4 *KTM*s. We have previously generated comparable H3K4me3 and H3K4me1 ChiP-seq data from neuronal nuclei obtained from the hippocampal CA region of mutant mice that lack either *Kmt2a* or *Kmt2b* from excitatory forebrain neurons (Kerimoglu *et al.*, 2017b). To ensure reliable comparison we reanalyzed in parallel the H3K4me3 and H3K4me1 Chip-seq datasets obtained from hippocampal neuronal nuclei of *Kmt2a*, *Kmt2b* and *Set1b* cKO mice. In line with the previous findings, all 3 *Kmt* mutant mice exhibit a substantial amount of TSS regions with deceased H3K4me3 **(Fig 4A)**. Interestingly, we also detected TSS regions with decreased H3K4me1 in all mutant mice, but only in *Setd1b* cKO mice a substantial number of TSS regions exhibited increased H3K4me1 **(Fig 4B)**. In addition, there was little overlap amongst the TSS regions with decreased H3K4me3 in *Kmt2a*, *Kmt2b* and *Setd1b* mutant mice, providing further evidence that *Setd1b* controls a unique gene-expression program in neuronal cells **(Fig 4C)**. To test this hypothesis directly, we decided to compare the corresponding gene-expression changes in the hippocampal CA1 region of the 3 KMT mutant mice. While the ChIP-seq data available for *Kmt2a* and *Kmt2b* mutant mice had been generated from neuronal nuclei, the corresponding gene-expression analysis represents RNA-seq data obtained from bulk tissue of the hippocampal CA1 region (Kerimoglu *et al.*, 2017b). To allow optimal comparison of these RNA-seq data to the gene-expression changes in *Setd1b* mutant mice, we also performed bulk RNA-seq from the hippocampal CA1 region of *Setd1b* cKO mice and control littermates. We observed 485 genes that were significantly down-regulated when comparing control to *Setd1b* cKO mice **(Fig 4D)**. These genes largely overlapped with the down-regulated genes detected via neuronal specific RNA-seq in *Setd1b* cKO mice **(Fig S4)**. While the total number of genes differentially expressed in the hippocampal CA1 region of *Kmt2a*, *Kmt2b* and *Setd1b* cKO mice was comparable, there was little overlap amongst them **(Fig 4E)**. Recently, RNA-sequencing data was reported for mice that were heterozygous for *Setd1a*. Although these mutants were heterozygous constitutive knock out mice and furthermore cortical tissue was analyzed instead of the hippocampus (Mukai *et al.*, 2019), it is interesting to note that there was virtually no overlap regarding the genes down-regulated in *Setd1a* knock out mice, when compared to the data obtained from our *Setd1b* cKO mice **(Fig. S5)**. Further support for a specific role of *Setd1b* in neuronal genes expression was revealed by the finding that the genes down-regulated in *Setd1b* cKO mice exhibited a significant enrichment for neuronal identity genes, while this was not the case for genes down-regulated in *Kmt2a* or *Kmt2b* cKO mice **(Fig 4F)**. In line with these data we observed that genes down-regulated in *Kmt2a* or *Kmt2B* cKO mice display decreased H3K4me3 at the TSS, while the levels of H3K4me1 were unaffected **(Fig 4G)**. In striking contrast, only the genes down-regulated in *Setd1b* cKO mice were characterized by reduced H3K4me3 and also increased H3K4me1 **(Fig 4G)**. Consequently, the genes that exhibit decreased H3K4me3 levels and were down-regulated in *Setd1b* cKO mice displayed significantly broader H3K4me3 peak at the TSS **(Fig 4H)** and were expressed a higher levels under basal conditions when compared the genes controlled by KMT2A or KMT2B **(Fig 4I)**. Functional pathway analysis showed that genes affected in the 3 KMT mutant mice represent different functional pathways. Interestingly, when compared to the gene-expression data obtained from *Kmt2a* or *Kmt2b* cKO mice, genes down-regulated in *Setd1b* cKO mice represent pathways intimately linked to learning and memory and the function of hippocampal neurons **(Fig 4J)**. Further analysis revealed that the genes affected in *Kmt2a* cKO mice are enriched for more general cellular processes, for processes related to gene-expression control and also neuronal plasticity related functions **(Fig S6A)**while the genes decreased in *Kmt2b* cKO mice represent almost exclusively pathways important for basal cellular function but not specifically important for neurons **(Fig S6B)**. In sum, these data support the view that *Setd1b* is of particular importance for the expression of genes essential for the identity of hippocampal neurons and synaptic plasticity. This view is further supported by the direct comparison of hippocampus-dependent memory function in *Setd1b*, *Kmt2a* and *Kmt2b* cKO mice. While loss of any of the 3 KMT’s leads to impaired spatial reference memory in the Morris water maze task (Kerimoglu *et al.*, 2013; Kerimoglu *et al.*, 2017b) (see Fig 1), memory impairment is more pronounced in *Setd1b* cKO **(Fig S7)**.

**Figure 4.**
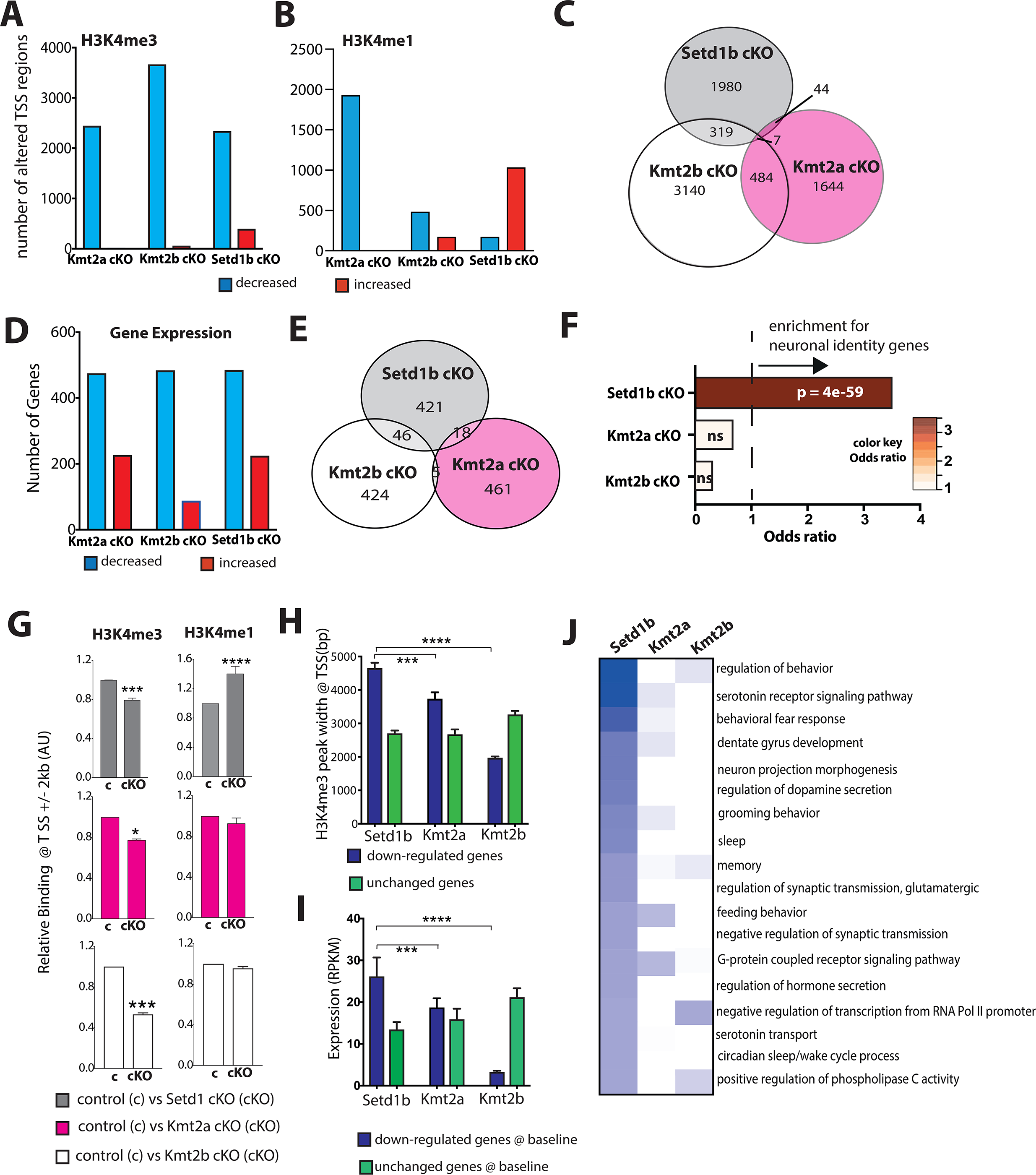
Comparative analysis of the hippocampal transcriptome in *Setd1b*, *Kmt2a* and *Kmt2b* cKO mice. **A.** Bar chart showing the number of genes that exhibit significantly altered H3K4me3 at the TSS (*Kmt2a*: control, n = 5; cKO, n = 3. *Kmt2b*: control, n = 6; cKO, n = 5. Setd1b: control, n = 4; cKO, n = 4). **B.** Bar chart showing the number of genes that exhibit significantly altered H3K4me1 at the TSS. **C.** Venn diagram comparing the genes with significantly decreased H3K4me3 at the TSS amongst in the 3 respective cKO mice. **D.** Bar chart showing the number of differentially expressed genes from bulk RNA-seq in each of the 3 KMT cKO mice. *Kmt2a*: control, n = 5; cKO, n = 6. *Kmt2b*: control, n = 8; cKO, n = 11. *Setd1b*: control, n = 6; cKO, n = 6) **E.** Venn diagram comparing the significantly down-regulated genes H3K4me3 amongst the 3 respective cKO mice. **F.** Donw-regualted genes with decreased H3K4me3 in each of the 3 KMT cKO mice were tested for the overlap to the 836 neuronal identity genes we had defined for the hippocampal CA region (See Fig S2). Only for *Setd1b* cKO mice a highly significant odds ratio (Fisher’s exact test) was observed, while there was no significant association amongst neuronal identity genes and the genes affected in *Kmt2a* and *Kmt2b* cKO mice. **G.** Left panel: Bar graphs showing H3K4me3 binding around the TSS of downregulated genes exhibiting significantly decreased H3K4me3 in either of the 3 KMT cKO mice (Two-way ANOVA: * *p*-value < 0.05, *** *p*-value < 0.001). Right panel depicts H3K4me1 for the same TSS regions (Two-way ANOVA **** *p*-value < 0.0001). Note that only in *Setd1b* cKO mice decreased H3K4me3 is accompanied by significantly increased H3K4me1. **H.** Genes exhibiting decreased H3K4me3 and reduced expression in *Kmt2a*, *Kmt2b* or *Setd1b* cKO mice were analyzed for H3K4me3 peak-width at the TSS under basal conditions. Genes affected in *Setd1b* cKO mice displayed significantly broader H3K4me3 peak-width when compared to genes down-regulated in *Kmt2a* or *Kmt2b* cKO mice. H3K4me3 peak-width at unchanged genes are shown for comparison. **I.** Bar graphs showing average basal expression of genes down-regulated with decreased H3K4me3 levels at the TSS in *Kmt2a*, *Kmt2b* or *Setd1b* cKO mice. Genes affected in *Setd1cKO* mice are expressed at significantly higher levels und basal conditions when compared to genes affected in *Kmt2a* or *Kmt2b* cKO mice. **J.**Heat map showing functional pathways of genes affected in *Kmt2a*, *Kmt2b* or *Setd1b* cKO mice. Note that genes affected by loss of *Setd1b* specifically represent pathways linked to neuronal function. Error bars indicate SEM.

### Single cell expression pattern likely contributes to the distinct role of Setd1b on neuronal gene-expression

In our effort to further elucidate the specific role of *Setd1b* on neuronal gene-expression and memory function we noticed that the levels of the H3K4 methyltransferases differ substantially in hippocampal neurons of the adult mouse brain. Surprisingly, our RNA-seq data from neuronal nuclei revealed *Setd1b* as the least expressed H3K4 methyltransferases when compared to *Kmt2a* or *Kmt2b* **(Fig 5A)**. These data might indicate that *Sedt1b* is generally expressed at very low levels or that alternatively only few cells may express *Setd1b*, a question that cannot be addressed on the basis our neuron-specific bulk RNA-seq. Thus, we decided to perform single nuclei sequencing. We isolated the hippocampus from 3-month old wild type mice and sorted NeuN + nuclei using our established protocol **(Fig 5B)**. These nuclei were then subjected to sequencing. As expected, we detected excitatory neurons of the cornu ammonis (CA) and dentate gyrus region as well as inhibitory neurons **(Fig 5C)**. Since our analysis so far was focused on the hippocampal CA region of mice that lack *Setd1b* from excitatory neurons, we selected the CA excitatory neurons and plotted the expression of *Kmt2a*, *Kmt2b* and *Setd1b*. In line with the data obtained from bulk sequencing of hippocampal neuronal nuclei, we observed that *Kmt2a* expression was most prominent when compared to *Kmt2b* or *Setd1b* **(Fig 5D)**. However, this difference was not due to the absolute expression value per cell, which was comparable for all 3 KMT’s **(Fig 5E)**. Rather, we observed that *Kmt2a* was expressed in the majority of the analyzed CA excitatory neurons, while *Kmt2b* and *Setd1b* were expressed in a comparatively small subset of cells **(Fig 5F)**. Next, we compared the gene-expression in neurons that either express *Kmt2a*, *Kmt2b* or *Setd1b* to cells that do not express the corresponding gene **(Fig 5E)**. A comparison of *Kmt2a* (+) vs *Kmt2a* (−) cells revealed that 897 genes were significantly increased in *Kmt2a* (+) cells, while 432 genes were expressed at higher levels in *Kmt2b* (+) and 214 genes in *Setd1b* (+) cells **(Fig 5G)**. Although the presence of *Kmt2a* affected more genes when compared to *Kmt2b* and *Setd1b,* the impact on gene-expression was comparatively small, which is indicated by the corresponding fold change of significantly differentially expressed genes. In fact, although the presence of *Setd1b* affected the least genes, the observed fold change of these genes was significantly greater when compared to *Kmt2* and *Kmt2b* **(Fig 5G)**. Only 7% of the genes significantly increased in *Kmt2a* (+) vs *Kmt2a* (−) cells showed a fold change greater than 1.5 **(Fig 5G)**. In case of *Kmt2b* (+) cells 41% of the regulated genes showed a fold change greater than 1.5 and for *Setd1b* this was true for 66 % of the regulated genes **(Fig 5G)**. These data suggest that the presence of *Kmt2a* contributes to the expression of many genes while its impact seems to be limited, when compared for example to the impact that the presence of *Setd1b* has on gene-expression. Hence, *Setd1b* affects comparatively less genes but with greater impact. When we subjected the genes specifically enriched in *Kmt2a* (+), *Kmt2b* (+) or *Setd1b* (+) cells to gene ontology and pathways analysis we observed that genes increased in the presence of *Setd1b* at the single cell level represent pathways linked to hippocampal function and interestingly also histone-acetylation **(Fig 5H)**, while these pathways were much less affected in *Kmt2a* (+) and *Kmt2b* (+) cells **(Fig. 5H)**. This is line with the finding that also at the single cell level, there is little overlap between the genes specifically enriched in either *Kmt2a* (+), *Kmt2b* (+) or *Setd1b* (+) cells **(Fig 5G.)**. We also calculated the eigen-value of the genes increased in *Setd1b* (+) cells and analyzed its expression *Setd1b* (+) and (−) cells as well as in cells positive for *Kmt2a* or *Kmt2b* **(Fig 5I)**. In line with our data, the eigen-value of *Setd1b*-specific genes was significantly increased in *Set1b* (+) cells compared to *Setd1b* (−) cells but also in cells positive for *Kmt2a* or *Kmt2b* **(Fig5I).** In summary, these data further support a specific role of Setd1b in neuronal function and suggest that hippocampal neurons expressing *Setd1b* in addition to other H3K4 KMTs may have a “plasticity benefit”.

**Figure 5.**
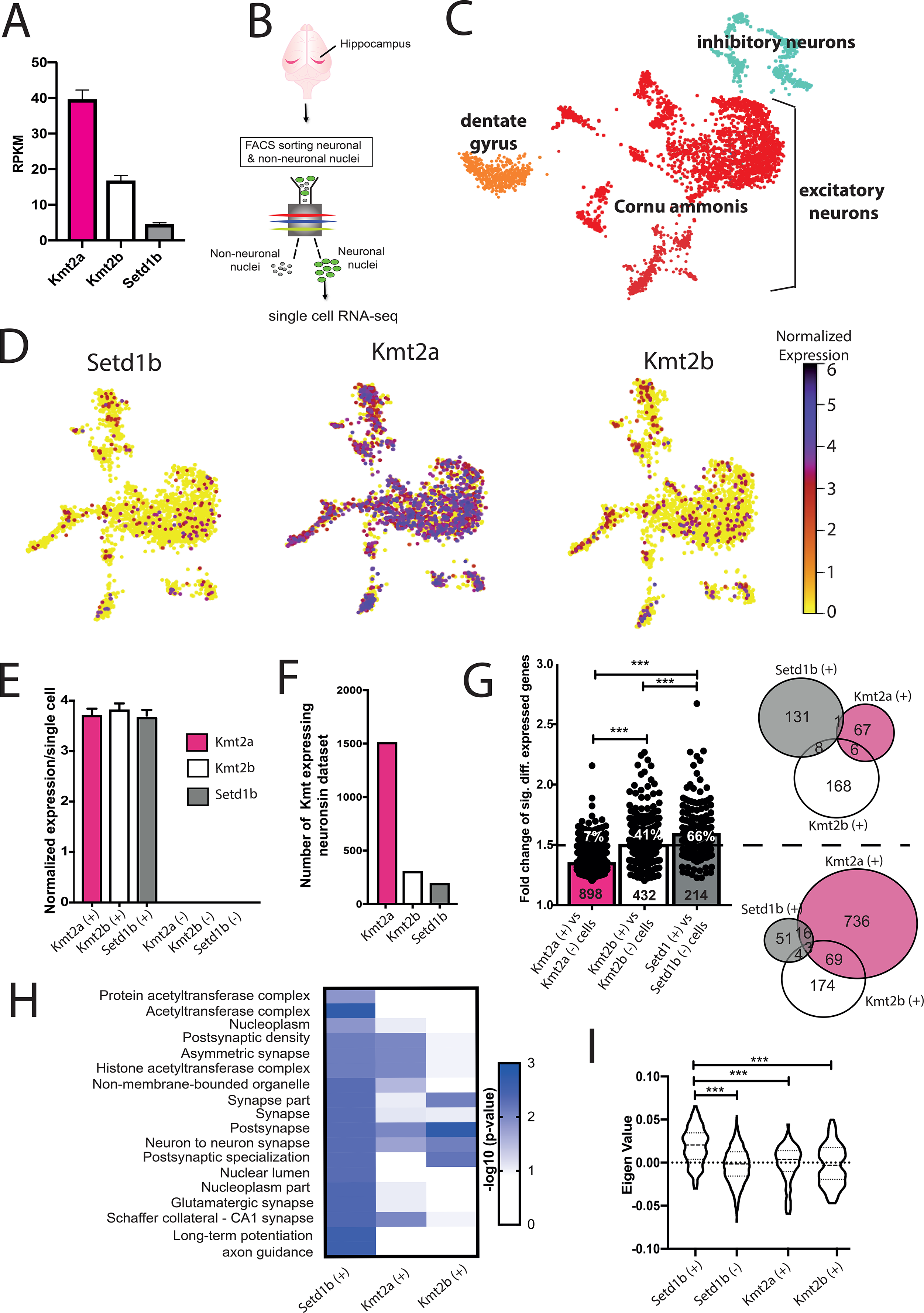
*Kmt2*, *Kmt2b* and *Setd1b* expression at the single cell level reveals a specific role for *Setd1b*. **A.** Bar graph showing the expression of *Kmt2*, *Kmt2b* and *Setd1b* expression in neuronal nuclei form the hippocampal CA region n = 3798. **B.** Experimental scheme for the single nuclei RNAseq experiment. **C.** UMAP plot showing the data from 3798 neuronal nuclei. **D.** UMAP plot showing the clustering of 2619 nuclei from hippocampal excitatory CA neurons indicating the normalized expression of *Setd1b* (left panel), *Kmt2a* (middle panel) and *Kmt2b* (right panel). **E.** Bar graph showing the normalized expression of *Kmt2*, *Kmt2b* and *Setd1b* in the respective positive cells. Note that the absolute expression amongst the 3 KTMs in the respective positive cells is not different. **F.** Number of nuclei positive for of the *Kmt2a*, *Kmt2b* or *Setd1b* in our dataset. **G.** We performed a differential expression analysis for *Kmt2a*, *Kmt2b* and *Setd1b* positive nuclei vs. the nuclei that did not express the corresponding KMT. Left panel: The bar graph shows the fold change of genes significantly increased in either *Kmt2a*, *Kmt2b* or *Setd1b* positive nuclei. The number in the bars refer to the number of differentially expressed genes. Please note that the majority of the genes significantly enriched in *Kmt2a* (+) cells exhibit a rather low fold change, while this is the opposite for *Setd1b* (+) cells. White number within the individual plotted genes indicate the percentage of genes that are significantly increased in *Kmt2a*, *Kmt2b* or *Setd1b* positive nuclei by a fold change greater than 1.5. Right panel: Venn diagram comparing the genes significantly increased in *Kmt2a*, *Kmt2b* or *Setd1b* positive nuclei with a fold change above 1.5 (upper diagram) or below a fold change of 1.5 (lower diagram). **H.** Top GO and Kegg pathways representing the genes increased in *Setd1b* positive nuclei. For comparison the enrichment of the same GO-terms/pathways is shown for genes enriched in *Kmt2a* or *Kmt2b* positive nuclei. **I.**Violin plot showing the eigen-value of the gene significantly altered when comparing *Setd1b* (+) to *Setd1b* (−) cells in *Setd1b* (+), *Setd1b* (−), *Kmt2a (+) and Kmt2b* (+) cells. Please note that this *Setd1b* specific gene-set is also significantly higher expressed when compared to *Kmt2a* or *Kmt2b* positive cells (One-way ANOVA *P* < 0.0001; *F* = 61.62; asterisks indicate unpaired t-Test; \*\*\**P* < 0.001)

## DISCUSSION

We show that loss of *Setd1b* from excitatory forebrain neurons impairs learning and memory in mice. These data are in line with previous findings showing that hippocampal H3K4me3 increases in response to memory training in rodents (Gupta *et al.*, 2010), while its levels are reduced in the hippocampus of a mouse for AD-like neurodegeneration (Gjoneska *et al*, 2015b)and in postmortem human brain samples of patients suffering from cognitive diseases (Shulha *et al.*, 2012). Our data furthermore support previous genetic studies linking mutations in *Setd1b* to intellectual disability (Hiraide *et al.*, 2018) (Labonne *et al.*, 2016). Since in our work *Setd1b* is not deleted during brain development but only in the postnatal brain, the presented findings suggest that reduced *Setd1b* expression can lead to cognitive dysfunction independent of developmental alterations. Impaired hippocampus-dependent memory has been also observed in mice that lack *Kmt2a* (Gupta *et al.*, 2010; Kerimoglu *et al.*, 2017b) or *Kmt2b* (Kerimoglu *et al.*, 2013)from excitatory neurons of the adult forebrain. Interestingly, mice heterozygous for *Setd1a*, the close homologue to *Setd1b* that is genetically linked to schizophrenia, show no impairment in the water maze task but rather exhibit impaired working memory and schizophrenia-like phenotypes (Mukai *et al.*, 2019). These data suggest that the different H3K4 KMT’s, at least *Kmt2a*, *Kmt2b*, *Setd1a* and *Setd1b*, serve distinct functions in the adult brain. The molecular characterization of *Setd1b* cKO further confirms this view. In line with the role of *Setd1b* in regulating H3K4me4, we observed a substantial decrease of neuronal H3K4me3 and the vast majority of these changes were observed at the TSS region of genes. Our data furthermore revealed that many genes with decreased H3K4me3 also exhibited reduced H3K9ac, which is in line with previous data showing that H3K4me3 appears to be a pre-requisite for H3K9ac, most likely since H3K4 KMT’s interact with histone-acetyltransferases (Kerimoglu *et al.*, 2013; Wang *et al*, 2009). For example, both *Setd1b* and the histone-acetlytransferase *Kat2a* where shown to interact with WDR5 (Lin *et al*, 2016; Ma *et al*, 2018), which is interesting since loss of *Kat2a* from excitatory forebrain neurons also leads to severely impairment of spatial reference memory (Stilling *et al.*, 2014). Somewhat unexpected was the observation that H3K4me1 levels were increased at a substantial number of TSS regions that exhibited decreased H3K4me3. A similar observation has however been made in yeast that expresses only one H3K4 KMTs, namely *Set1* (Soares *et l.*, 2017). The authors show that highly expressed genes have the broadest H3K4me3 peak at the TSS, while moderate to low expressed genes are characterized by a narrow H3K4me3 peak and comparatively higher H3K4me1 levels. This H3K4me-dependent pattern of gene-expression was directly correlated to degree of SET1 activity at the TSS. In agreement with these data we observed that the genes with decreased H3K4me3 and increased H3K4me1 in *Setd1b* cKO mice are indeed characterized by a broad H3K4me3 peak at the TSS and high expression levels at the basal state. It is in this context interesting to note that *Setd1b* is the closest mammalian orthologue of the yeast SET1 protein, from which the other H3K4 methyltransferases have evolved (Shilatifard, 2012). Importantly, when we analyzed gene-expression in *Setd1b* cKO mice we observed that not all genes that exhibit reduced H3K4me3 display reduced mRNA expression. Rather, we found that specifically genes with decreased H3K4me3 and increased H3K4me1, hence only the genes with the broadest H3K4me3 distribution at the TSS were significantly down-regulated in *Setd1b* cKO mice. These genes represent key pathways linked to memory formation and the identity of hippocampal neurons, which is also in line with a recent study reporting that memory training specifically activates hippocampal genes with broad H3K4me3 peaks at the TSS (Collins *et al*, 2019). In sum, these data suggest that *Setd1b* is important for the expression of neuronal identity genes in the hippocampus, that are linked to learning and memory processes, a function that might be specifically associated with *Setd1b*. Thus, when we directly compared genes down-regulated in *Setd1b*, *Kmt2a* or *Kmt2b* cKO mice, the genes down-regulated in *Setd1b* cKO mice were characterized by significantly broader H3K4me3 peaks at the TSS, significantly higher baseline expression and they were enriched for neuronal identity genes and pathways specifically important for learning-related processes. A significant gradient *Setd1b* > *Kmt2a* > *Kmt2b* was observed for all of these comparisons. In line with these data, also the memory performance in the water maze training was more severely affected in *Setd1b* cKO mice followed by mice lacking *Kmt2a* or *Kmt2b*. These data may suggest that H3K4 KTM’s other than *Setd1b* are essential to ensure the sufficient expression of genes important for basal cellular processes in neurons, while *Setd1* enables to preeminent expression of neuronal identity genes. This view is in line with a recent study in mouse embryonic stem cells in which *Setd1b* was associated with the expression of highly expressed genes that exhibit a broad H3K4me3 peak, while *Kmt2b* was linked to the expression of genes with narrow H3K4me3 peaks (Sze *et al*, 2020). Interestingly, this study suggested a functional redundancy of *Setd1b* and *Setd1a*. It is likely that this is true for gene-expression programs related to more general cellular processes. However, the situation might be different for genes specifically important in post-mitotic neurons. Moreover, mutations in either *Setd1a* or *Setd1b* lead to distinct neuropsychiatric diseases and unlike Setd1b cKO mice, *Setd1a* heterozygous mutant mice do not exhibit impairment of long-term memory consolidation (Mukai *et al.*, 2019). At present we cannot conclusively answer the question how *Setd1b* affects the expression of specifically neuronal identity genes. Previous data suggest that H3K4 KMT’s associate with different co-activators (Lee *et al*, 2006), (Dreijerink *et al*, 2006; Shilatifard, 2012) (Hughes *et al*, 2004; Yokoyama *et al*, 2004) (Scacheri *et al*, 2006) which could explain the regulation of specific gene-expression programs. Our data provides a complimentary explanation that should be considered in addition. Using single-nucleus-sequencing we observed that *Setd1b* is expressed in a comparatively small number of hippocampal neurons, when compared for example to *Kmt2a*. Yet, the impact on memory function is most significant when hippocampal neurons lack *Setd1b*, as for example compared to the more abundantly expressed *Kmt2a* or *Kmt2b*. The comparison of *Setd1b* expressing hippocampal neurons to cells that do not express *Setd1b* confirmed a specific role for *Setd1b* in the expression of genes intimately linked to neuronal function and memory processes. This was different for *Kmt2a* and *Kmt2b* expressing neurons suggesting that the presence of *Setd1b* at the single cell level enables particularly efficient expression of genes important for the function of hippocampal neurons and memory consolidation. It has to be re-iterated that these genes are also detectable in neurons that lack *Setd1b* and express *Kmt2a* or *Kmt2b,* but to a lesser extent (See Fig 5I). This allows for some interesting hypotheses. For example, a number of studies demonstrated that upon learning a specific set of neurons – in most cases neurons that initiate a *cFos*-dependent gene-expression program – become part of a neuronal circuitry important for memory encoding. Particularly important are those neurons that are later reactivated during memory retrieval (Reijmers *et al*, 2007; Tonegawa *et al*, 2018) (Josselyn SA, 2019). All of these studies found that only a small fraction of the originally activated cells became reactivated during memory retrieval. It is thus tempting to speculate that the activity of genes such as *Setd1b* might help to shape the neuronal ensemble that will indeed be reactivated during memory retrieval, a hypothesis that would need to be tested in further studies. Taking into account that decreased neuronal H3K4me3 levels have been observed in cognitive and neurodegenerative diseases therapeutic strategies that reinstate specifically the expression of neuronal plasticity genes controlled by *Setd1b* might be particularly helpful. We suggest that the various epigenetic drugs currently tested in pre-clinical and clinical settings for cognitive diseases should especially be analyzed for their potential to reinstate the H3K4me3 peak width at neuronal identity genes.

In conclusion, we show that *Setd1b* is essential for memory consolidation and ensures the proper expression of neuronal identity genes. Since *Setd1b* is expressed only in a subset of hippocampal neurons it may provide a plasticity benefit to those cells thereby regulating memory formation at the molecular level. In turn, *Setd1b*-related gene-expression programs could be a suitable therapeutic target to treat cognitive diseases and help patients suffering from intellectual disability.

## MATERIALS AND METHODS

### Animals

All animals used in this study were C57BL/6J mice and of 3-6 months of age. The experimental groups were age and sex matched. Mice were kept in standard home cages with food and water provided *ad libitum*. All experiments were performed according to the animal protection law of the state of Lower Saxony.

### Behavior experiments

The behavioral experiments were performed as described previously(Kerimoglu *et al.*, 2017b). For in depth feature analysis from water maze data, a modified version of MUST-C algorithm was used (Illouz *et al.*, 2016).

### Tissue isolation and processing

Hippocampal CA tissues were dissected from WT and *Setd1b* cKO mice, flash frozen in liquid nitrogen and stored at −80°C until further processing.

### Cell-type specific nuclear RNA isolation and sequencing

Frozen CA tissues from left and right hemisphere of two mice were pooled together and processed on ice to maintain high RNA integrity. Tissue was homogenized using a plastic pestle in a 1.5mL Eppendorf tube containing 500 uL EZ prep lysis buffer (Sigma, NUC101-1KT) with 30 strokes. The homogenate was transferred into 2 mL microfuge tubes, lysis buffer was added up to 2 mL and incubated on ice for 7 minutes. After centrifuging for 5 minutes at 500g supernatant was removed and the nuclear pellet was resuspended into 2 mL lysis buffer and incubated again on ice (7 minutes). After centrifuging for 5 minutes at 500g, the supernatant was removed and the nuclei pellet was resuspended into 500ul nuclei storage buffer (NSB: 1x PBS; Invitrogen, 0.5% RNase free BSA;Serva, 1:200 RNaseIN plus inhibitor; Promega, 1x EDTA-free protease inhibitor; Roche) and filtered through 40 μm filter (BD falcon) with additional 100 μL NSB to collect residual nuclei from the filter.

Nuclei were stained with anti-NeuN-Alexa488 conjugated antibody (1:1000) for 45 minutes and washed once with NSB. Stained nuclei were then FACS-sorted with FACSaria III using 85 μm nozzle. Nuclei were gated by their size, excluding doublets and neuronal nuclei were separated from non-neuronal nuclei by their NeuN-Alexa488 fluorescence signal. Sorted nuclei were collected into a 15 mL falcon tube precoated with NSB, spun down and RNA was isolated using Trizol LS. After addition of chloroform according to the Trizol LS protocol, aqueous phase was collected and RNA was isolated by using Zymo RNA clean & concentrator-5 kit with DNAse treatment. Resulting RNA concentration were measured in Qubit and RNA-seq was performed using 100ng of neuronal RNA with illumina TruSeq RNA Library Prep Kit. Since glial nuclei are smaller and contains very little amount of RNA, neuronal nuclear RNA was scaled down and 1ng from both neuronal and glial nuclear RNA was used to make RNA-seq libraries using Takara SMART-Seq v4 Ultra Low Input RNA Kit. Libraries were sequenced using single-end 75 bp in Nextseq 550 or single-end 50 bp in HiSeq 2000, respectively.

### Cell-type specific chromatin isolation and ChIP sequencing

Frozen tissues were homogenized, formaldehyde (1%) fixed for 10 minutes and quenched with 125mM glycine for 5 minutes. Debris was removed by sucrose gradient centrifugation. The resulting nuclear pellet was stained with anti-NeuN-Alexa488 conjugated antibody (1:1000) for 25 minutes and washed 3 times with PBS. Stained nuclei were then FACS sorted with FACSaria III using 85 μm nozzle. Nuclei were gated similarly as described previously(Halder *et al*, 2016). Sorted nuclei were collected into a 15mL falcon tube and transferred into 1.5mL tubes. The nuclear pellet was flash frozen in liquid nitrogen and saved at −80°C for further processing. For chromatin shearing, the pellet was resuspended into 100uL RIPA buffer (containing 1% SDS) and sonicated for 25 cycles in Diagenode bioruptor plus with high power and 30 cycles on/ 30 cycles off. Chromatin shearing was checked by taking a small aliquot and decrosslinking the DNA by 30 minutes RNAse and 2 hours of proteinase K treatment. DNA was isolated using SureClean Plus protocol. Sheared chromatin size was determined using Bioanalyzer 2100(DNA high sensitivity kit) and the concentration was measured using Qubit 2.0 fluorometer (DNA high sensitivity kit). 0.3μg chromatin was used along with 1 μg of antibody to do ChIP for H3K4me3 (Abcam ab8580), H3K4me1 (Abcam ab8895), H3K27ac (Abcam ab4729) and H3K9ac (Millipore 07-352). ChIP was performed as previously described(Halder *et al.*, 2016). The resulting ChIP DNA was subjected to library preparation using NEBNext Ultra II DNA library preparation kit and sequenced for single end 50bp at illumina HiSeq 2000.

### ChIP-Seq Analysis

Base calling and fastq conversion were performed using Illumina pipeline. Quality control was performed using fastqc (www.bioinformatics.babraham.ac.uk/projects/fastqc). Reads were mapped to mm10 mouse reference genome with STAR aligner v2.3.0.w. PCR duplicates were removed by *rmdup-s* function of samtools. BAM files with unique reads belonging to the same group were merged into a single BAM file with the *merge* function of samtools. Profile plots were created from these merged BAM files with NGSPlot. Peak calling was performed using MACS2 against the input corresponding to the particular group (i.e., control or cKO) using q < 0.1. Consensus peaksets were generated for each histone modification individually using the Diffbind package of Bioconductor with the command *dba.count* and the parameter *minOverlap=1*. Then, these consensus peaksets were intersected with each other using the *intersect* function of bedtools with default parameters.. The differential binding analysis for each histone mark between control and *Setd1b* cKO was then performed using Diffbind with this common peakset as input. For the comparison of H3K4me3 and H3K4me1 changes in *Kmt2a* cKO, *Kmt2b* cKO and *Setd1b* cKO common peaksets for each individual histone mark from three separate ChIP-Seq experiments were extracted. In this case, first, consensus peaksets for a histone mark from each individual ChIP-Seq experiment (i.e., “Control vs Kmt2a cKO”, “Control vs Kmt2b cKO” and “Control vs Setd1b cKO”) were determined using Diffbind. For the purpose of comparing the effects of the three KMT knockdowns on H3K4me3 or H3K4me1 the differential binding analyses for each individual ChIP-Seq experiment were performed always utilizing these common consensus peaksets. Diffbind package was used for differential binding analysis with in-built DESEQ2 option for differential analysis. The annotation of the genomic regions was performed with HOMER.

### RNA-Seq Analysis

Base calling, fastq conversion, quality control, mapping of reads to mouse reference genome (mm10) were performed as described before (Kerimoglu *et al.*, 2017b)Seq. Reads were counted using FeaturesCount (http://bioinf.wehi.edu.au/featureCounts/). Differential expression was analyzed with DESeq2 package of Bioconductor RPKM values were calculated using edgeR package of Bioconductor.

### Single-nucleus RNA-Seq

Unfixed NeuN+ neuronal nuclei were isolated as mentioned above (section: Cell-type specific nuclear RNA isolation and sequencing). Sorted neuronal nuclei were counted in a Neubauer chamber with 10% trypan blue (in PBS) and nuclei concentration were adjusted to 1000 nuclei/μL. The nuclei were further diluted to capture and barcode 4000 nuclei according to Chromium single cell 3ʹ reagent kit v3 (10X genomics). Single nuclei barcoding, GEM formation, reverse transcription, cDNA synthesis and library preparation were performed according to 10X genomics guidelines. Finally, the library was sequenced in Illumina NextSeq 550 according to manufacturer’s protocol. Gene counts were obtained by aligning reads to the mm10 genome (GRCm38.p4)(NCBI:GCA_000001635.6) using CellRanger software (v.3.0.2) (10XGenomics). The CellRanger count pipeline was used to generate a gene-count matrix by mapping reads to the pre-mRNA as reference to account for unspliced nuclear transcripts. The dataset contained 3841 cells with a mean of 31.053 total read counts over protein-coding genes.

The SCANPY package was used for pre-filtering, normalization and clustering (Wolf *et al*, 2018) Initially, cells that reflected low-quality cells (either too many or too few reads, cells isolated almost exclusively, cells expressing less than 10% of house-keeping genes (Eisenberg & Levanon, 2013) were excluded remaining in 3801 cells. Next, counts were scaled by the total library size multiplied by 10.000, and transformed to log space. A total of 3066 highly variable genes were identified based on dispersion and mean, the technical influence of the total number of counts was regressed out, and the values were rescaled. Principal component analysis (PCA) was performed on the variable genes, and UMAP was run on the top 50 principal components (PCs) (Becht *et al*, 2018), The top 50 PCs were used to build a k-nearest-neighbours cell–cell graph with k= 200 neighbours. Subsequently, spectral decomposition over the graph was performed with 50 components, and the Louvain graph-clustering algorithm was applied to identify cell clusters. We confirmed that the number of PCs captures almost all the variance of the data. For each cluster, we assigned a cell-type label using manual evaluation of gene expression for sets of known marker genes. Two cell-type clusters identified as neurons from dentate gyrus and inhibitory neurons were excluded. Remaining excitatory neuronal cells from CA region were re-clustered using the same settings as described above. For each cluster, differentially expressed genes were detected using the Wilcoxon rank-sum test as implemented in the function rank_genes_groups in SCANPY.

## Data availability

All RNA and ChIP-seq datasets will be made available via GEO database

## Acknowledgments

This work was supported by the following third party funds to AF: The ERC consolidator grant DEPICODE (648898), the BMBF projects ENERGI (01GQ1421A) and Intergrament (01ZX1314D), and funds from the German Center for Neurodegenerative Diseases FS was supported by the DFG grant SA1005/2-1and funds from the DZNE. AM is a student of the International Max Planck Research School (IMPRS) for Genome Science. SS& RI are students of the IMPRS for Neuroscience.

## Author contributions

AM initiated the project as part of her PhD thesis, performed behavior experiments, performed and analyzed immunohistochemistry and analyzed mutant mice; SS generated cell-type specific ChIP/RNA-seq and single nucleus RNA-seq data and contributed to the analysis of cell type specific RNA-seq, CK analyzed and interpreted ChIP-seq and RNA-seq data and supervised all bioinformatic data analysis, DMF analyzed single nucleus RNA-seq data, LK, XX & EMZ helped with the generation of single nucleus RNA-seq; AK & FAS provided material and analyzed data, AM, SS, CK and AF designed experiments and generated figures. CK and AF wrote the paper.

---

**Expanded view Fig 1.**
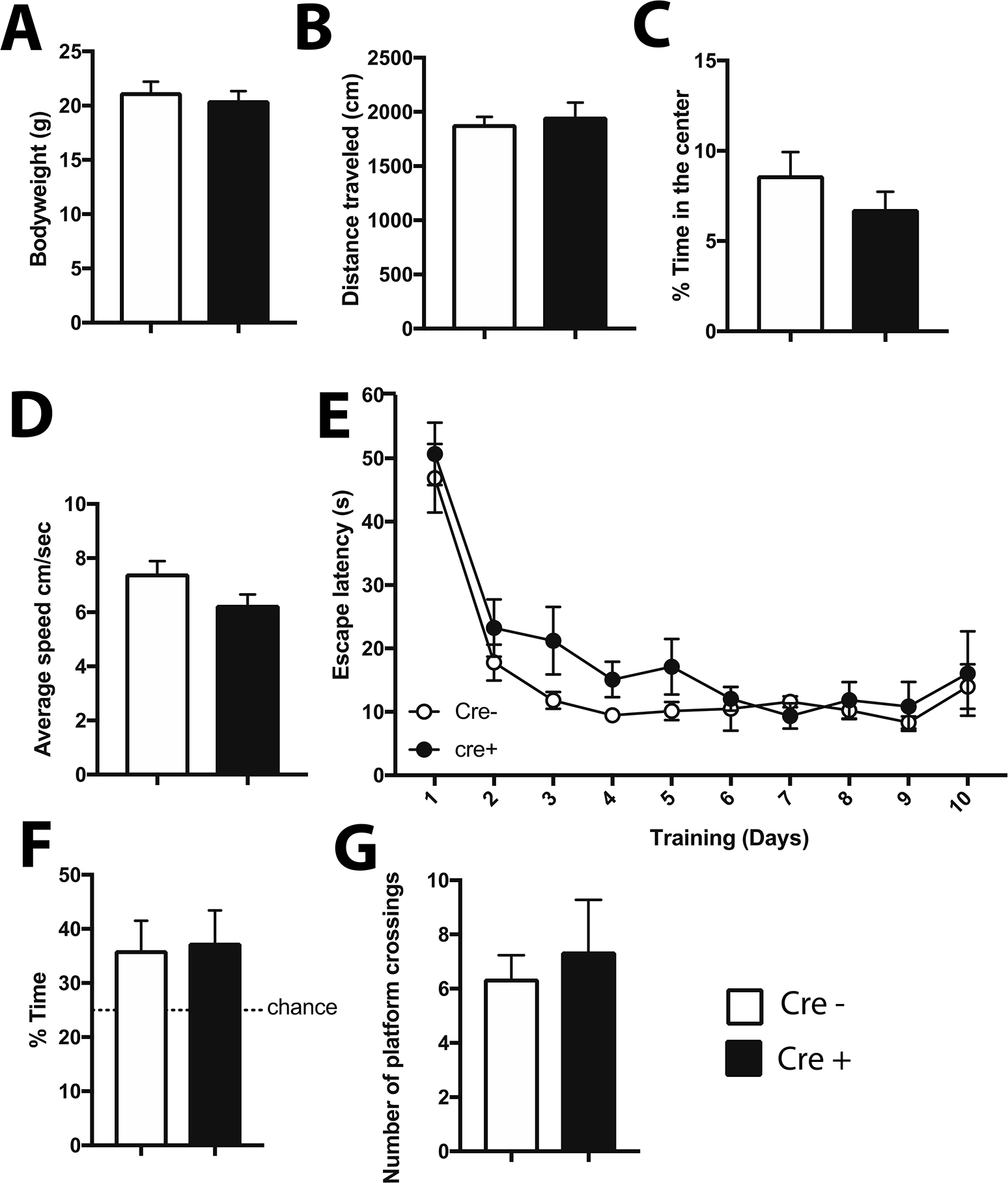
Behavioral analysis of mice expressing CamKII-driven Cre recombinase. **A.** Transgenic mice expressing CRE under control of the CamKII promoter were subjected to behavior testing (n=8, Cre +) comparing them to wild type mice from the same breeding colony that did not express CRE (n=8; Cre -). No difference was observed in body weight. **B.** The distance traveled in the open filed test and **(C)** the time spent in the center of the arena was similar amongst groups. **D.** No difference in the swimming speed was observed amongst groups when subjected to the water maze test. **E.** Escape latency during water maze training was similar in CRE - and CRE + mice. **F.** During the probe test performed after 10 trainings days, CRE - and CRE + mice showed similar performance then time spent in the target quadrant and **(G)** the number of platform crossings were analyzed. Error bars indicate SEM.

**Expanded view Fig 2:**
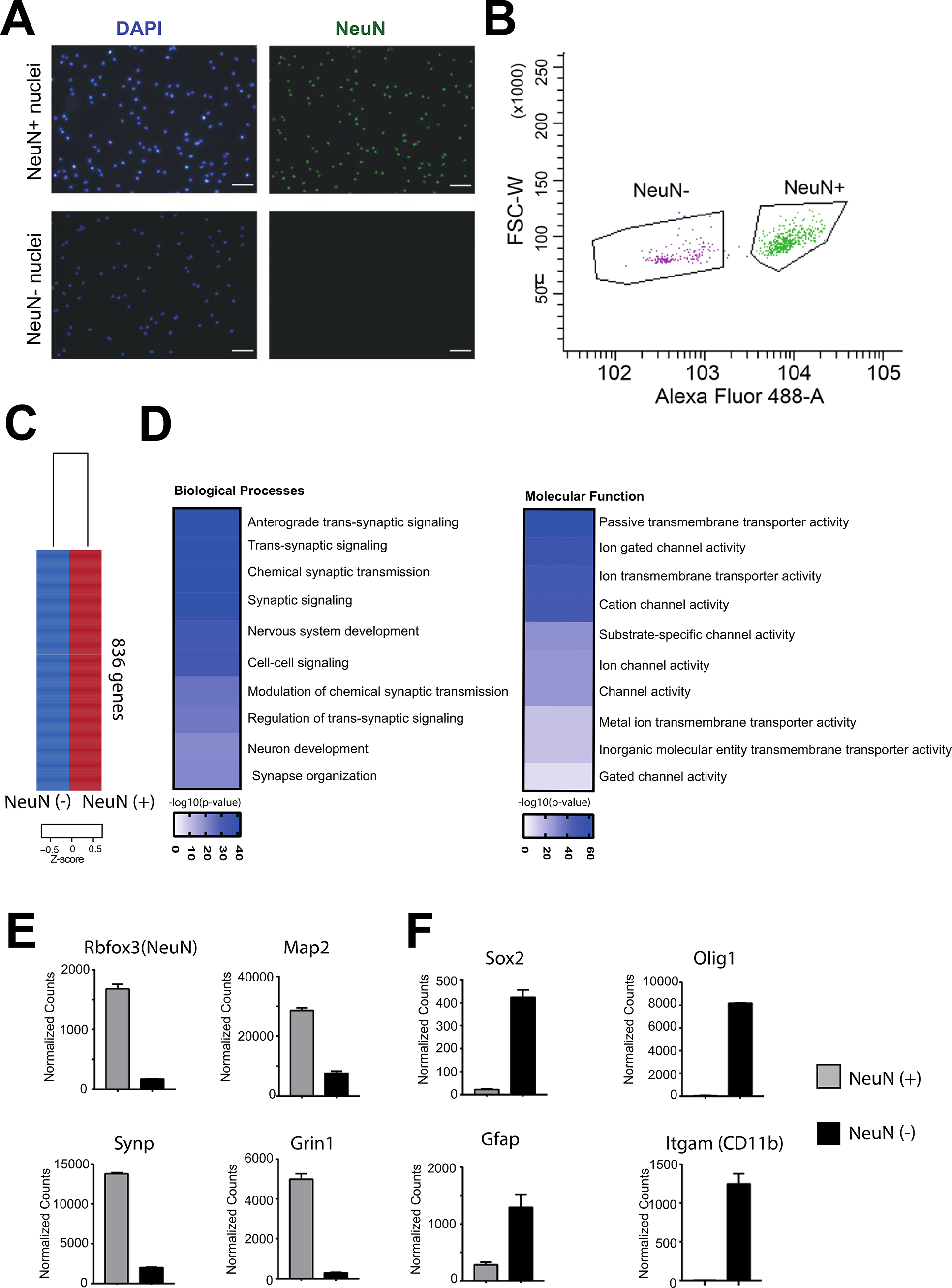
Sorting neuronal and non-neuronal nuclei for RNA-analysis. Nuclei from the hippocampal CA region were subjected to FACS as depicted in Fig 2A. **A**. Representative images showing nuclei that were sorted using the neuronal marker NeuN. Note that no NeuN positive nuclei are detected in the NeuN (−) fraction confirming the puirity of the approach. Scale bar: 50μm **C.** Gating strategy for NeuN (+) and NeuN (−) nuclei sorting. **C**. RNA-sequencing (n=2/group) was performed from NeuN (+) and NeuN (−) nuclei and a differential expression analysis was performed. Heat map shows 836 genes specifically enriched in NeuN (+) nuclei when compared to NeuN (−) nuclei. The criteria to select those genes were: adjusted p value <0.01, basemean >150, fold change > 5. **D.** GO-term analysis showing that the top 10 enriched biological processes and molecular functions for the 836 genes enriched in NeuN (+) nuclei all represent specific neuronal processes. **E.** Normalized expression values obtained from the RNA-seq experiment showing the expression of selected genes known to be enriched in neurons. **F.** Normalized expression values of genes that are known to be enriched in non-neuronal cells including glia cells. Error bars indicate SEM.

**Expanded view Fig. 3.**
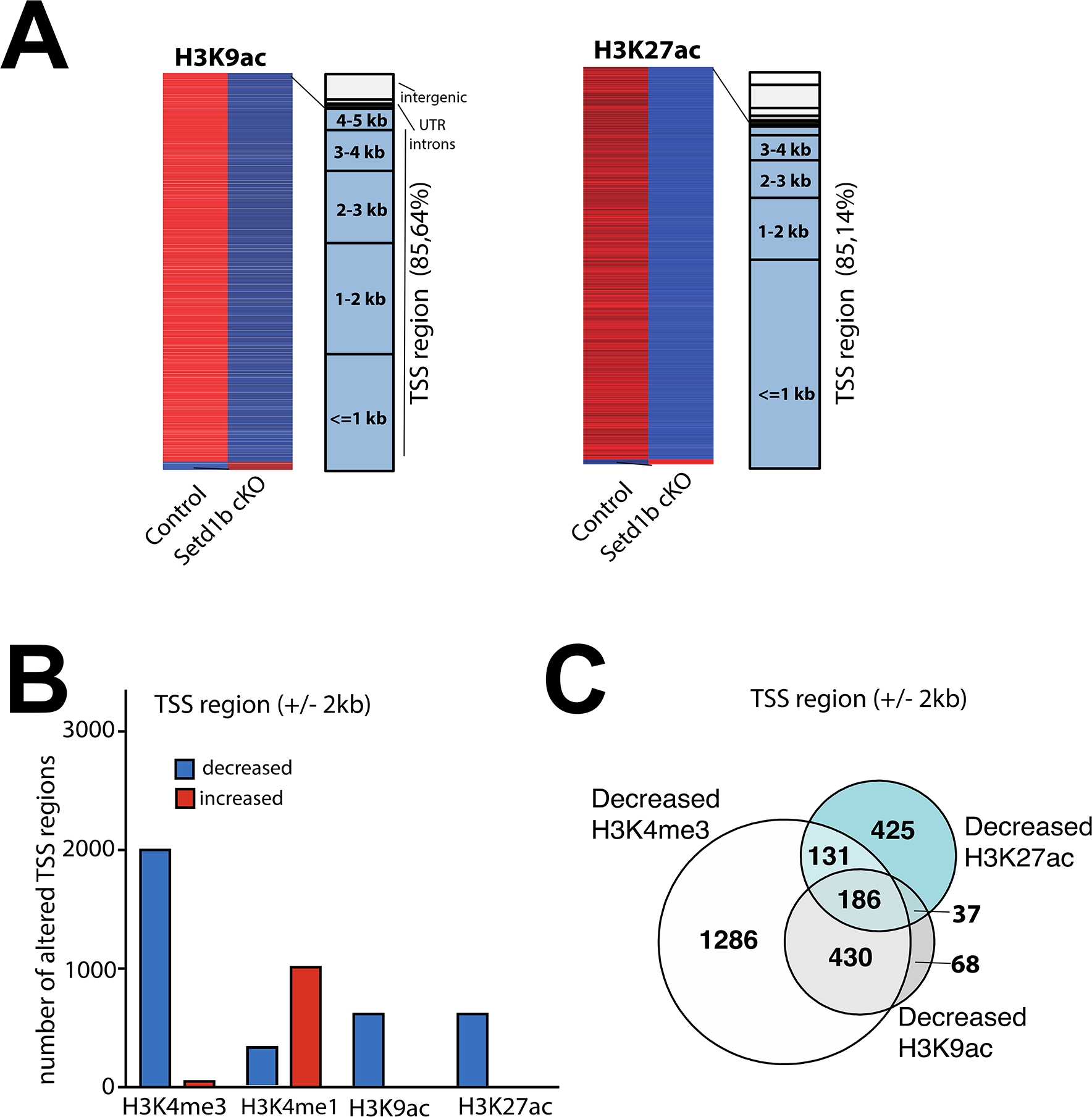
Decreased H3K9ac and H3K27ac in *Setd1b* cKO mice. **A.** Left panel: Heat map showing genes with differential H3K9ac sites at the TSS in neuronal nuclei from *Setd1b* cKO mice and their genomic location. Right panel shows the same analysis for H3K27ac. **B.** Bar chart showing the number of genes with decreased and increased H3K9ac and H3K27ac marks at the TSS region. Data for H3K4me3 and H3K4me1 are shown for comparison. As expected, the most affected histone-mark is H3K4me3. **C.** Venn diagram showing that most of the sites exhibiting decreased H3K9ac at the TSS also exhibit reduced H3K4me3, while this was not the case for H3K27ac. TSS, transcription start site.

**Expanded view Fig 4.**
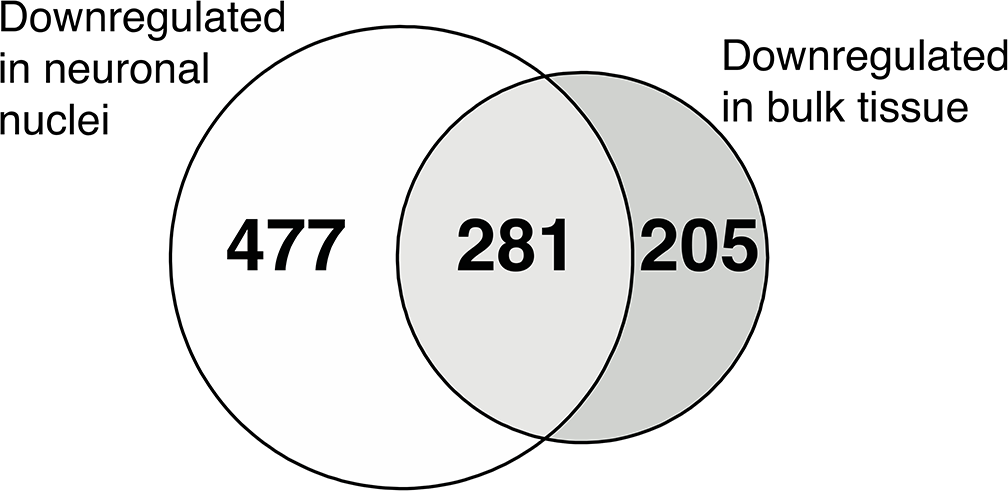
Comparison of gene-expression changes in *Setd1b* cKO mice detected from cell-type specific and bulk tissue RNA-seq. Venn diagram showing the overlap of genes down-regulated in *Setd1b* cKO mice detected via RNA-seq from neuronal nuclei or bulk hippocampal CA tissue. Please note that more genes are detected when neuronal nuclei are analyzed suggesting that some of the difference are masked by cell type heterogeneity when bulk tissue is analyzed.

**Expanded view Fig. 5.**
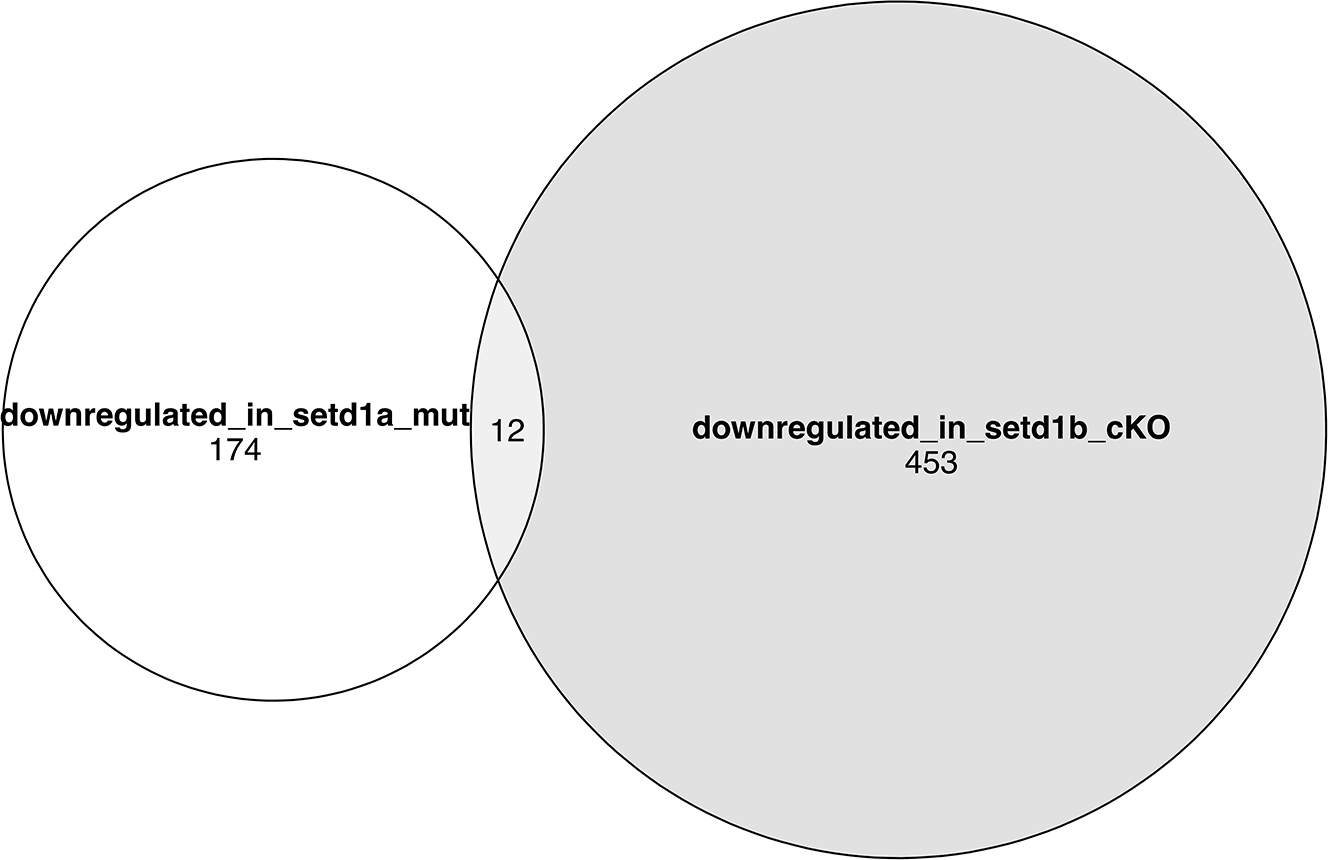
Comparison of the genes down-regulated in *Setd1A* and *Setd1b* mutant mice. Venn diagram showing genes down-regulated in *Setd1A* vs *Setd1b* cKO mice. Please note that genes affected in the different mutant mice are very different. Of course, care has to be taken since the data from *Setd1A* mutant mice stems from a recent publication by Mukai et al., 2019 (PMID:31606247). In this study cortical tissue from heterozygous mice constitutively lacking *Setd1A* were analyzed, while our data stems from the hippocampus of conditional knock out mice.

**Expanded view Fig. 6.**
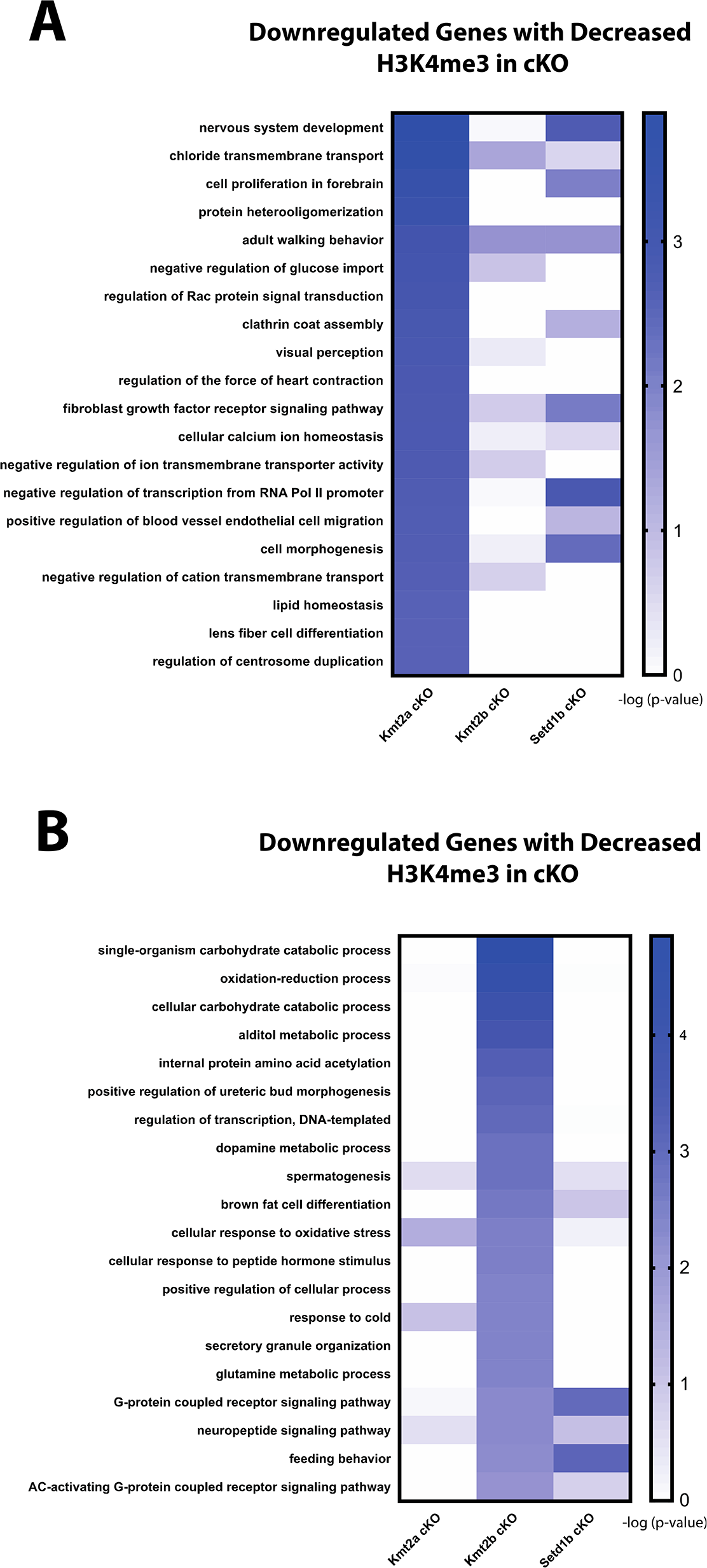
Functional pathways affected in *Kmt2a* and *Kmt2b* cKO mice. **A.** Heat map showing functional pathways analysis for genes down-regulated in *Kmt2a* cKO mice. Enrichment of the same pathways is also shown for *Kmt2b* and *Setd1b* cKO mice. **B.** Heat map showing functional pathways analysis for genes down-regulated in *Kmt2b* cKO mice. Enrichment of the same pathways is also show for *Kmt2a* and *Setd1b* cKO mice. Please note that the pathways affected in *Kmt2a* or *Kmt2b* cKO mice differ substantially from those affected in *Setd1b* cKO mice. All data is based on comparable RNA-seq data generated from bulk hippocampal CA1 region.

**Expanded view Fig. 7.**
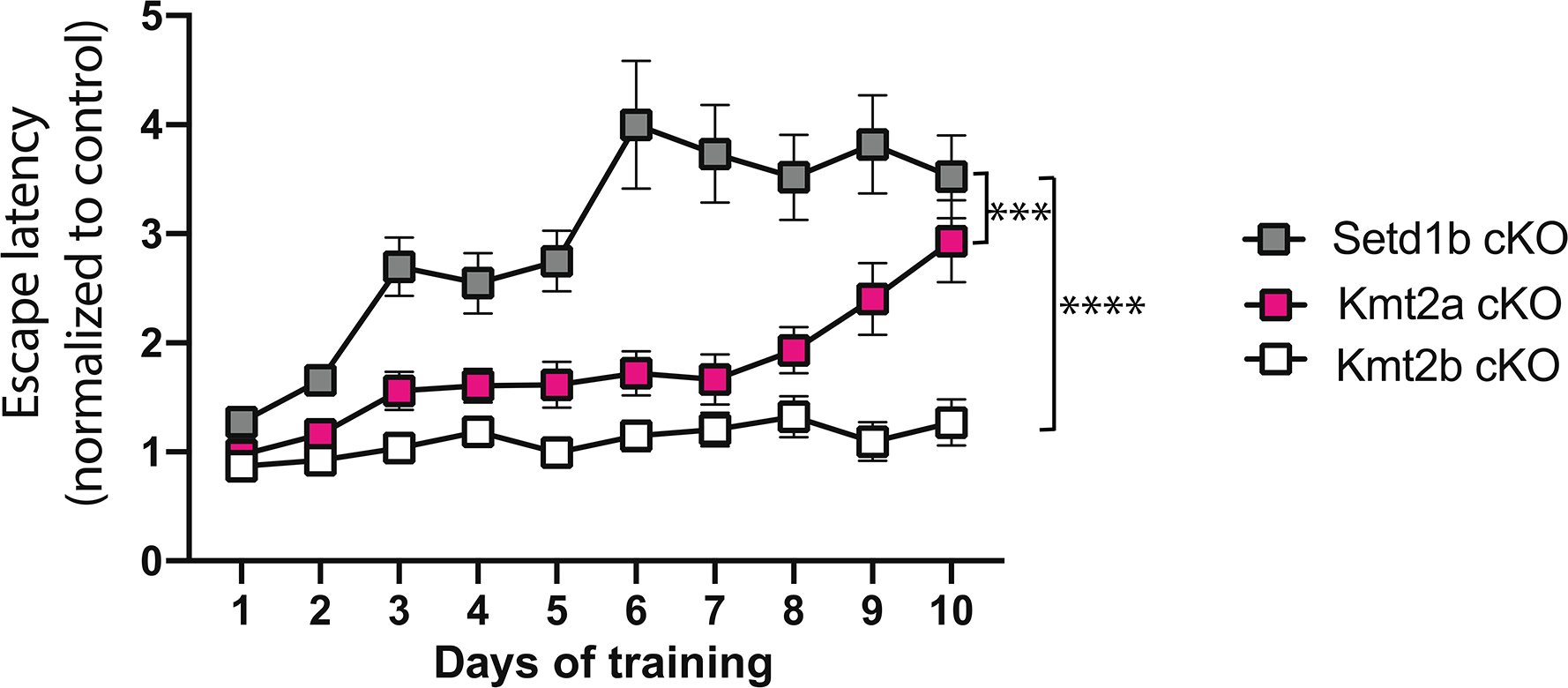
Spatial reference learning in *Kmt2a*, *Kmt2b* and *Setd1b* cKO mice. To compare spatial reference learning in the 3 different mutant mice, we normalized the data to the corresponding control group. This is important, since the experiments were performed at different time points. The data on *Kmt2a* and *Kmt2b* cKO mice was generated in our laboratory and is already published (Kerimoglu *et al.*, 2013) (Kerimoglu *et al.*, 2017a). The data on *Setd1b* cKO mice were generated as part of this project using the same protocol. In this plot an increase in the normalized escape latency shows the difference to the corresponding control group. Hence, a higher normalized escape latency indicates a greater difference to the corresponding control and this more severe learning impairment. This difference in normalized escape latency is significantly greater in *Setd1b* cKO mice when compared to *Kmt2a* or *Kmt2b* cKO mice. It is interesting to note that the degree of memory impairment seems to parallel the gene-expression data, hence *Setd1* appears to be most important for the expression of neuronal identity genes linked to learning and memory *Setd1b* cKO mice are also most affected in spatial memory formation. Loss of *Kmt2a* affects some pathways specific to neuronal function but also more general cellular processes while loss of *Kmt2b* seem to affect genes that are not specific to neuronal function. In line with this, loss of Kmt2b has the least effect on spatial memory and in fact memory defects in *Kmt2b* cKO mice only become obvious during prolonged training and the probe test^23^. *Setd1b* cKO (n = 14) vs *Kmt2a* cKO (n = 13): Repeated measures ANOVA, genotype effect: F (1,25) = 16.83, *** *p*-value < 0.001. *Setd1b* cKO (n = 14) vs *Kmt2b* cKO (n = 22): Repeated measures ANOVA, genotype effect: F (1,34) = 70.66, **** *p*-value < 0.0001.

